# Male-specific vasotocin expression in the medaka tuberal hypothalamus: androgen dependence and probable role in aggression

**DOI:** 10.1101/2023.09.13.557517

**Authors:** Yukika Kawabata-Sakata, Shinji Kanda, Kataaki Okubo

## Abstract

Terrestrial vertebrates have a population of androgen-dependent vasotocin (VT)-expressing neurons in the extended amygdala that are more abundant in males and mediate male-typical social behaviors, including aggression. Teleosts lack these neurons but instead have novel male-specific VT-expressing neurons in the tuberal hypothalamus. Here we found in medaka that *vt* expression in these neurons is dependent on post-pubertal gonadal androgens and that androgens can act on these neurons to directly stimulate *vt* transcription via the androgen receptor subtype Ara. Furthermore, administration of exogenous VT induced aggression in females and alterations in the androgen milieu led to correlated changes in the levels of tuberal hypothalamic *vt* expression and aggression in both sexes. However, genetic ablation of *vt* failed to prevent androgen-induced aggression in females. Collectively, our results demonstrate a marked androgen dependence of male-specific *vt* expression in the teleost tuberal hypothalamus, although its relevance to male-typical aggression needs to be further validated.

## Introduction

Aggression is an adaptive behavioral trait that is crucial for competition for territories, food, and mating partners and the establishment of social hierarchies. Although the regulation of aggression in vertebrates involves many different neural mechanisms, particular attention has been given to two hormonal systems in the brain: sex steroids and nonapeptides (Kelly and Wilson, 2020).

Sex steroids are peripherally derived or produced in the brain and act on neural circuits to modulate behavior, primarily through binding to specific nuclear receptors that serve as ligand-gated transcription factors (Yang and Shah, 2014). Androgens, among other sex steroids, play a central role in facilitating aggression, and males are typically more aggressive than females due to their androgen-dominated steroid milieu (Hashikawa *et al*., 2018; Lischinsky and Lin, 2020). In rodents, the stimulatory effects of androgens on aggression are largely mediated by the activation of estrogen receptors (ESRs) after their conversion to estrogens in the brain (Yang and Shah, 2014). Recent studies have revealed that activation of the ESR subtype ESR1 in the ventromedial hypothalamus (VMH) is particularly important for the expression of aggressive behavior (Chen and Hong, 2018; Hashikawa *et al*., 2018; Lischinsky and Lin, 2020). However, the transcriptional targets of ESR1 that mediate aggressive behavior remain elusive, and the specific role of sex steroids in the regulation of aggression is unclear. Furthermore, and importantly, it is unlikely that the findings in rodents apply to other vertebrates including primates and teleost fish, where androgens act directly on behaviorally relevant neural circuits via the androgen receptor (AR) without conversion to estrogens (Okubo *et al*., 2019; 2022).

Nonapeptides, namely, vasotocin (VT, also called vasopressin in mammals) and oxytocin (OT), are evolutionarily conserved neuropeptides that have been associated with a wide range of social behaviors, including aggression (Theofanopoulou *et al*., 2021; Mennigen *et al*., 2022). The largest population of neurons expressing VT and that expressing OT both lie in the paraventricular nucleus (PVN), where they project throughout the brain, as well as to the pituitary, to modulate behavior (Rigney *et al*., 2022). In many terrestrial vertebrates, additional neuronal populations expressing VT in an androgen-dependent, and hence male-biased, manner have been identified in the extended amygdala, specifically in the bed nucleus of the stria terminalis (BNST) and the medial amygdala (MeA) (Kelly and Goodson, 2014; Aspesi and Choleris, 2022; Rigney *et al*., 2022; 2023). These neurons serve as the main regulators of male-typical social behaviors, and elicit pro- or anti-aggressive behavioral responses in males, depending on species and social context (Aspesi and Choleris, 2022).

Notably, however, teleost fish lack VT-expressing neurons in the extended amygdala, even though their aggression—like that of terrestrial vertebrates—seems to depend on VT and is generally more prevalent in males (Godwin and Thompson, 2012; Rigney *et al*., 2023). This suggests that, in teleosts, VT may function within another neural circuit to elicit high levels of aggression in males. In line with this idea, teleosts have several populations of VT-expressing neurons in the tuberal hypothalamus, in addition to the major population that spans the brain nucleus homologous to the PVN and its immediate surroundings (Godwin and Thompson, 2012; Oldfield *et al*., 2015). Our previous findings further revealed that, in medaka fish (*Oryzias latipes*), the populations in the posterior tuberal nucleus (NPT) and the posterior part of the ventral tuberal nucleus (pNVT) are confined to males (Kawabata *et al*., 2012). The NPT and pNVT are considered homologous to the ventral tegmental area/substantia nigra and the anterior hypothalamus, respectively (Forlano and Bass, 2011; Loveland and Hu, 2018), both of which have been implicated in nonapeptide-regulated social behavior (Rigney *et al*., 2022).

To our knowledge, no information is available on the regulation or role of VT-expressing neuronal populations in the NPT and pNVT, but it has been reported in pupfish (*Cyprinodon nevadensis amargosae*) that VT expression in the hypothalamus is higher in socially dominant, highly aggressive males (Lema *et al*., 2015). Taken together, the above observations led us to hypothesize that VT expression in either or both of these neuronal populations is induced exclusively in males in an androgen-dependent manner and contributes to the high levels of aggression typical of males. Here we tested this hypothesis by investigating the regulatory mechanisms and physiological roles of male-specific VT expression in the tuberal hypothalamus of medaka.

## Materials and Methods

### Animals

All experimental procedures involving animals were performed in accordance with the University of Tokyo Institutional Animal Care and Use Committee guidelines. The committee requests the submission of an animal-use protocol only for use of mammals, birds, and reptiles, in accordance with the Fundamental Guidelines for Proper Conduct of Animal Experiment and Related Activities in Academic Research Institutions under the jurisdiction of the Ministry of Education, Culture, Sports, Science and Technology of Japan (Ministry of Education, Culture, Sports, Science and Technology, Notice No. 71; June 1, 2006). Accordingly, we did not submit an animal-use protocol for this study, which used only teleost fish and thus did not require approval by the committee.

Wild-type medaka of the d-rR strain and *vt* knockout medaka produced in this study were kept under controlled conditions at 28 °C and a photoperiod of 14:10 light/dark, and were fed with live *Artemia nauplii* and dry food (Otohime; Marubeni Nisshin Feed, Tokyo, Japan) 3–4 times a day. Sexually mature fish between 3 and 6 months of age were used in all experiments except for the analysis of age-dependent changes in *vt* expression, for which fish of 1, 2, 3, and 7 months of age were employed. To control for genetic diversity and environmental variation, siblings raised in the same conditions were assigned as the comparison group in all experiments, including those with knockout fish. All sampling was done 1–2.5 hours after initiation of the light period.

### Gonadectomy and drug treatment

A small incision was made in the ventrolateral abdominal wall of anesthetized fish (0.02% tricaine methane sulfonate). The gonad was removed through the incision, which was then closed with nylon thread. Following a 3-day recovery period in saline (0.9% sodium chloride), gonadectomized fish were reared for 6 days in water containing 100 ng/ml of 11-ketotestosterone (KT; the primary androgen in teleosts that cannot be converted to estrogens) (Cosmo Bio, Tokyo, Japan) or estradiol-17β (E2; the primary estrogen in teleosts and other vertebrates) (Fujifilm Wako Pure Chemical, Osaka, Japan), or vehicle alone (ethanol) and then sampled. Sham-operated control fish were subjected to the same surgical procedure as gonadectomized fish but without removing the gonad and then treated with vehicle alone.

In separate experiments, males and females (including *vt* knockout females) with intact gonads were reared for 9 days in water containing 250 ng/ml of the AR antagonist cyproterone acetate (CA) (LKT Laboratories, St. Paul, MN) and 100 ng/ml of KT, respectively. These fish were observed daily for changes in aggressive behavior and, when necessary, were sampled on day 9. The concentration of sex steroids used was determined based on steroid levels in medaka serum (Tilton *et al*., 2003). Ovarian-intact females were also treated intraperitoneally with 0.02 ng/mg body weight of Vt peptide (Fujifilm Wako Pure Chemical) or vehicle alone (saline) and observed for changes in aggressive behavior 2 hours after treatment.

### Single-label *in situ* hybridization

A digoxigenin (DIG)-labeled cRNA probe for *vt* was generated by PCR amplification of a DNA fragment corresponding to nucleotides 1–845 (845 bp) of the medaka *vt* cDNA followed by *in vitro* transcription using T7 RNA polymerase and DIG RNA Labeling Mix (Roche Diagnostics, Basel, Switzerland). The *in situ* hybridization procedure has been outlined in detail elsewhere (Hiraki-Kajiyama *et al*., 2019). Briefly, brains were fixed in 4% paraformaldehyde (PFA), paraffin embedded, and coronally sectioned at 10-μm thickness. Hybridization signal was visualized with an anti-DIG antibody conjugated to alkaline phosphatase (RRID: AB_514497; Roche Diagnostics) and nitro blue tetrazolium/5-bromo-4-chloro-3-indolyl phosphate (NBT/BCIP) substrate (Roche Diagnostics). The color was allowed to develop for 5 hours. All sections to be compared were processed in the same batch. To obtain quantitative data, all relevant sections were photographed and converted to black and white binary images by thresholding using Adobe Photoshop (Adobe, San Jose, CA), and the total area of *vt* expression signal across all the relevant sections was calculated for each brain nuclei using ImageJ (https://imagej.nih.gov/ij/).

### Double-label *in situ* hybridization

The double-label *in situ* hybridization procedure has been described in detail elsewhere (Kawabata-Sakata *et al*., 2020). Briefly, brains were fixed in 4% PFA, embedded in 20% sucrose/5% agarose, and cryosectioned at 20-μm thickness in the coronal plane. Sections were simultaneously hybridized with the *vt* probe, which was labeled with fluorescein using T7 RNA polymerase and Fluorescein RNA Labeling Mix (Roche Diagnostics), and a DIG-labeled AR (*ara*, NM_001122911; *arb*, NM_001104681) probe (Hiraki *et al*., 2012). Fluorescein was detected with an anti-fluorescein antibody conjugated to horseradish peroxidase (RRID: AB_2737388; PerkinElmer, Waltham, MA) and visualized using the TSA Plus Fluorescein System (PerkinElmer); DIG was detected with an anti-DIG antibody conjugated to alkaline phosphatase (RRID: AB_514497; Roche Diagnostics) and visualized with Fast Red (Roche Diagnostics). Sections were counterstained with 4′,6-diamidino-2-phenylindole (DAPI) to identify cell nuclei. Fluorescent images were captured using a TCS SP8 confocal laser-scanning microscope (Leica Microsystems, Wetzlar, Germany) with the following excitation/emission wavelengths: 405/410–480 nm (DAPI), 488/495–545 nm (fluorescein), and 552/565–700 nm (Fast Red).

### Transcriptional activity assay

A bacterial artificial chromosome (BAC) clone containing the medaka *vt* locus (clone ID: ola1-127B15) was obtained from National BioResource Project (NBRP) Medaka (http://www.shigen.nig.ac.jp/medaka/). The *vt* gene and flanking regions were sequenced and analyzed for the presence of potential canonical bipartite androgen-responsive element (ARE)-like sequences using Jaspar (version 5.0_alpha; http://jaspar.genereg.net/). cDNA fragments encoding full-length medaka Ara and Arb (NM_001104681 and NM_001170833, respectively) were amplified by PCR and inserted into the expression vector pcDNA3.1/V5-His-TOPO (Thermo Fisher Scientific, Waltham, MA). Fragments of genomic DNA upstream of the first methionine codon (2677 bp) and downstream of the stop codon (1469 bp) of *vt* were amplified by PCR from the BAC clone and inserted into the respective NheI and XbaI sites of the luciferase reporter vector pGL4.10 (Promega, Madison, WI). The resultant reporter construct was transiently transfected into COS-7 cells using Lipofectamine LTX (Thermo Fisher Scientific), together with an internal control vector pGL4.74 (Promega) and either the Ara or Arb expression construct. Six-hour post-transfected cells were treated with 0, 10^−10^, 10^−8^, or 10^−6^ M KT for 18 hours in Dulbecco’s modified Eagle’s medium (phenol red-free) containing 5% charcoal-treated fetal bovine serum (Thermo Fisher Scientific). After cell lysis, luciferase activity was determined using the Dual-Luciferase Reporter Assay System (Promega) on the GloMax 20/20n Luminometer (Promega). All assays were conducted in duplicate and repeated independently three times.

To determine the ARE responsible for androgen induction of *vt* transcription, assays were also conducted with luciferase reporter constructs containing only the fragment upstream of the first methionine codon or carrying point mutations in the identified ARE-like sequences. A construct containing the upstream fragment was prepared as described above. Constructs containing point-mutated ARE-like sequences were prepared using the PrimeSTAR Mutagenesis Basal Kit (Takara Bio, Shiga, Japan). Because an ARE half-site can function alone to confer androgen inducibility (Pihlajamaa *et al*., 2015), both half-sites of each ARE-like sequence were mutated (into a HindIII recognition site sequence [AAGCTT] to facilitate confirmation of the mutation). The procedures for cell transfection, KT treatment, and luciferase activity measurements were the same as above, except that a single dose (10^−6^ M) of KT was used.

Additional assays were performed with luciferase reporter constructs containing truncated versions of the downstream fragment to see the effects of simultaneous loss of multiple ARE-like sequences in this fragment. Constructs containing the full-length (1469 bp) or serially 3′-truncated (970 bp and 485 bp) downstream fragment were generated as described above and used in these assays. Cell transfection, KT treatment (10^−6^ M), and luciferase activity measurements were performed as described above.

### Aggressive behavior test

The aggressive behavior test was conducted as previously described (Yamashita *et al*., 2020). Briefly, four fish of the same genotype and sex that were not familiar with one another were housed in a 2-liter rectangular tank and separated from each other by opaque partitions. After 10-min acclimation to the tank, the partitions were removed to allow the fish to interact. Their behavior was recorded for 30 min with a digital video camera (Everio GZ-G5; Jvckenwood, Kanagawa, Japan, or iVIS HF S11/S21; Canon, Tokyo, Japan). All tests were done 1–5 hours after initiation of the light period. Video recordings were manually analyzed for the total number of each aggressive act (chase, fin display, circle, strike, and bite).

### Production of knockout medaka

The vt knockout medaka line was produced using transcription activator-like effector nuclease (TALEN) technology, targeting the sequence corresponding to the mature Vt peptide (Supplementary Fig. 1A), essentially as described (Takahashi et al., 2016). In brief, TALE repeat arrays were assembled using the Joung Lab REAL Assembly TALEN kit (Addgene 1000000017). Synthesized TALEN mRNAs were injected into the cytoplasm of one-cell stage embryos. Once these fish reached adulthood, they were outcrossed to wild-type fish and the resulting progeny were tested for target site mutations by T7 endonuclease I assay (Kim et al., 2009) and direct sequencing. A founder fish was identified that reproducibly produced progeny carrying a 10-bp deletion that introduced a frameshift into the mature Vt peptide. The progeny were intercrossed to obtain homozygous, heterozygous, and wild-type siblings. Both male and female homozygous fish were viable and displayed no obvious morphological or developmental defects. Each fish was genotyped by direct sequencing of the targeted locus.

Only female knockout fish were used in this study because we considered that the most direct approach to test our hypothesis would be to confirm that exogenous androgens induce aggression in wild-type females but not in *vt* knockout females. We considered males to be unsuitable for this experiment because they have high levels of endogenous androgens and would be less susceptible to exogenous androgens.

### Statistical analysis

All quantitative data were presented as mean ± standard error of the mean. On graphs, individual data points were also plotted to indicate the underlying distribution. All statistics were analyzed using GraphPad Prism (GraphPad Software, San Diego, CA). Data between two groups were compared using unpaired two-tailed Student’s *t*-test. When the F-test indicated that the variances between groups were significantly different, Welch’s correction was applied. To compare data among more than two groups, one-way analysis of variance (ANOVA) was performed, followed by either Dunnett’s (comparisons between control and experimental groups) or Bonferroni’s (comparisons among experimental groups) *post hoc* test. In cases where the Bartlett’s and Brown-Forsythe tests revealed significant differences in variances among groups, the data were subjected to log transformation to correct for heterogeneity of variance. When variances remained heterogeneous following transformation, Kruskal-Wallis test was utilized, followed by Dunn’s *post hoc* test. For analyses of age-dependent changes in *vt* expression and aggressive behavior in *vt*-deficient females treated with KT, two-way ANOVA was used to determine the effects of and interactions between age and sex and between genotype and KT treatment, respectively. If a significant interaction was detected, differences between groups were further analyzed by Bonferroni’s *post hoc* test. All data points were included in the analyses and no outliers were defined.

## Results

### Male-specific *vt* expression in the tuberal hypothalamic nucleus is dependent on post-pubertal gonadal androgens

First, we determined the spatiotemporal pattern of *vt* expression and related sex differences in the medaka brain. *In situ* hybridization analysis of brains from different ages showed that *vt* was expressed in the PMp/PPa/PMm/PMg (the latter two brain nuclei are homologous to the PVN) in the preoptic area and the SC/aNVT, NAT, NPT, and pNVT in the tuberal hypothalamus at all ages examined (Fig. 1A and B; see Supplementary Table 1 for abbreviations of medaka brain nuclei). Signal quantification revealed that expression in the PMp/PPa/PMm/PMg, shared by males and females, increased with age similarly in both sexes, but levels were slightly higher in males overall (main effect of age, *p* < 0.0001; main effect of sex, *p* = 0.0020; interaction between age and sex, *p* = 0.5837) (Fig. 1C). Expression in both the SC/aNVT and NAT peaked at 2 months of age, when secondary sexual characters were well-developed but no spawning had yet occurred (spawning began at 3 months); there were no significant sex differences except for a slight female bias in the SC/aNVT at 2 months of age (*p* = 0.016 between sexes) (Fig. 1C). Expression in the NPT and pNVT was male-specific at all ages examined, with significant sex differences detected after 2 months of age (*p* = 0.0005, 0.0003, and 0.0078 for NPT at 2, 3, and 7 months, respectively; *p* < 0.0001 for pNVT at 2, 3, and 7 months) (Fig. 1C).

**Fig. 1.**
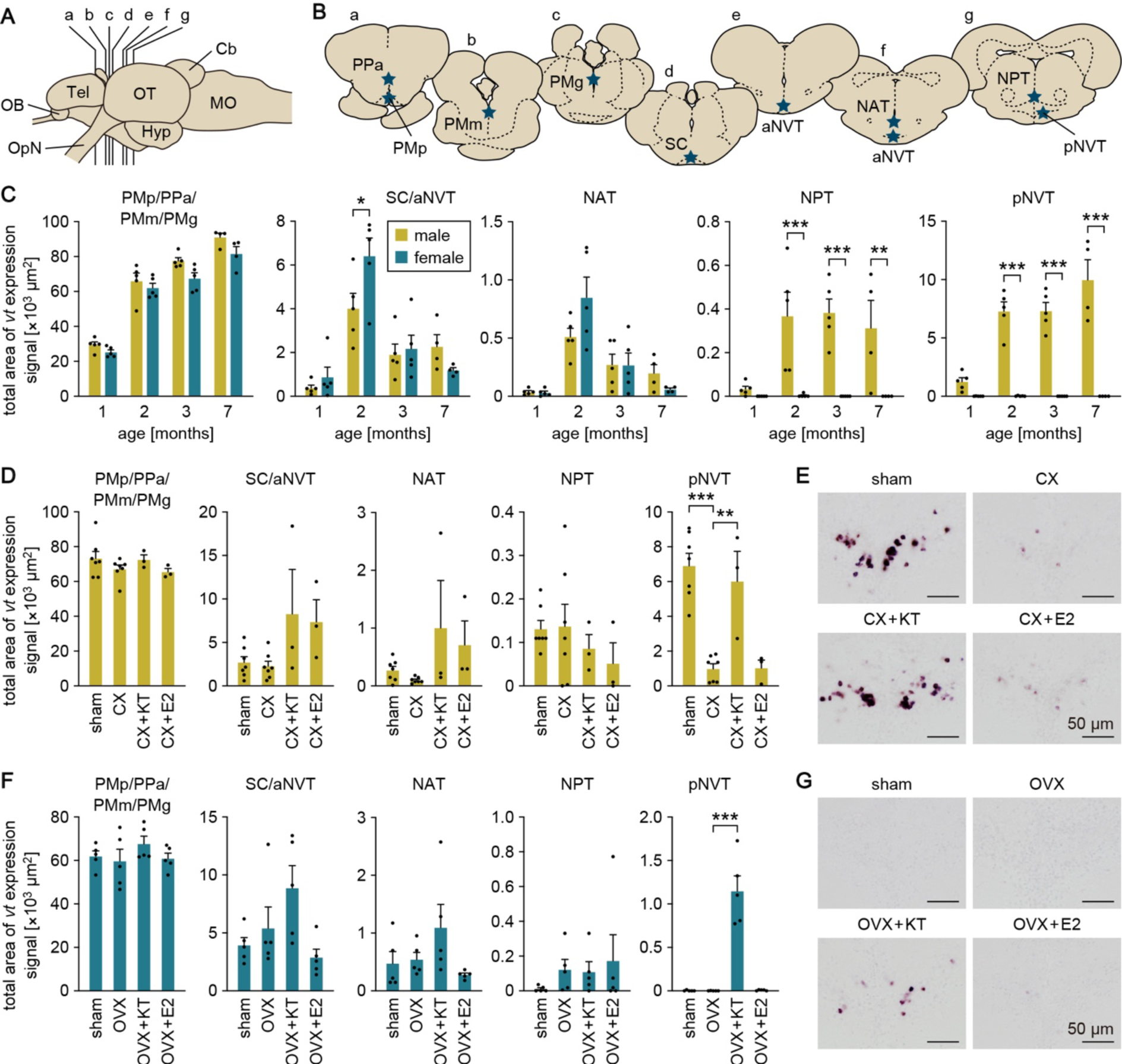
Male-specific *vt* expression in the tuberal hypothalamic nucleus is dependent on post-pubertal gonadal androgens. (A) Schematic drawing of the medaka brain (lateral view, anterior to left) depicting the approximate levels of sections shown in panel B. (B) Coronal brain sections containing nuclei in which *vt* is expressed (stars). See Supplementary Table 1 for abbreviations of brain regions and nuclei. (C) Total area of *vt* expression signal in each brain nucleus of males (yellow columns) and females (blue columns) at different ages (n = 5 per sex and age, except n = 4 at 7 months). (D) Total area of *vt* expression signal in each brain nucleus of sham-operated males (sham) and castrated males treated with vehicle alone (CX), KT (CX+KT), or E2 (CX+E2) (n = 7 for sham and CX; n = 3 for CX+KT and CX+E2). (E) Representative images of *vt* expression in the pNVT of sham, CX, CX+KT, and CX+E2 males. Scale bars are all 50 μm. (F) Total area of *vt* expression signal in each brain nucleus of sham females and ovariectomized females treated with vehicle alone (OVX), KT (OVX+KT), or E2 (OVX+E2) (n = 5 for each group). (G) Representative images of *vt* expression in the pNVT of sham, OVX, OVX+KT, and OVX+E2 females. Scale bars are all 50 μm. Statistical differences were determined by Bonferroni’s *post hoc* test (C, D, F). \**p* < 0.05; \*\**p* < 0.01; \*\*\**p* < 0.001.

These results led us to speculate that sex steroid hormones produced by the gonads after puberty onset influence the pattern of *vt* expression in the medaka brain. We tested this idea by quantitative *in situ* hybridization of *vt* expression in fish that were gonadectomized in adulthood and treated with KT or E2. In the male pNVT, castration significantly reduced *vt* expression (*p* < 0.0001), which was restored by KT treatment (*p* = 0.0014) but not by E2 treatment (Fig. 1D and E). In the female pNVT, *vt* expression was not detected under normal conditions but was induced by KT treatment following ovariectomy (*p* < 0.0001) (Fig. 1F and G). In other brain nuclei, no significant differences between treatments were observed (Fig. 1D and F). Collectively, these results suggest that high circulating levels of androgens released from the testis stimulate *vt* expression in the pNVT in post-pubertal males, whereas in females, the lack of androgen stimulation prevents its induction. In addition, the effect of androgens on *vt* expression in the pNVT is transient and reversible; thus, *vt* expression is attenuated in androgen-depleted adult males and, conversely, induced in estrogen-depleted and androgen-supplemented adult females.

### Androgens can directly activate the transcription of *vt* through Ara

To investigate the possible direct action of androgens on *vt*-expressing neurons in the pNVT, we first determined if these neurons coexpress ARs. Most teleosts, including medaka, possess two subtypes of AR, designated Ara and Arb (Okubo *et al*., 2022). Double-label *in situ* hybridization for *vt* and each AR subtype revealed that *ara*, but not *arb*, was abundantly expressed in the pNVT of both sexes, and virtually all of the *vt*-expressing neurons in the male pNVT were positive for *ara* expression (Fig. 2A and b). Note that the names of *ara* and *arb* used in the present study follow the nomenclature of Ogino *et al*. (2016; 2023), which reflects the orthology/paralogy relationships of teleost ARs. Thus, the gene referred to as *ara* in our previous publications (*e.g*., Hiraki *et al*., 2012; 2014; Kawabata-Sakata *et al*., 2020; Yamashita *et al*., 2017; 2020) is denoted as *arb* in this study, and the gene referred to as *arb* as *ara*.

**Fig. 2.**
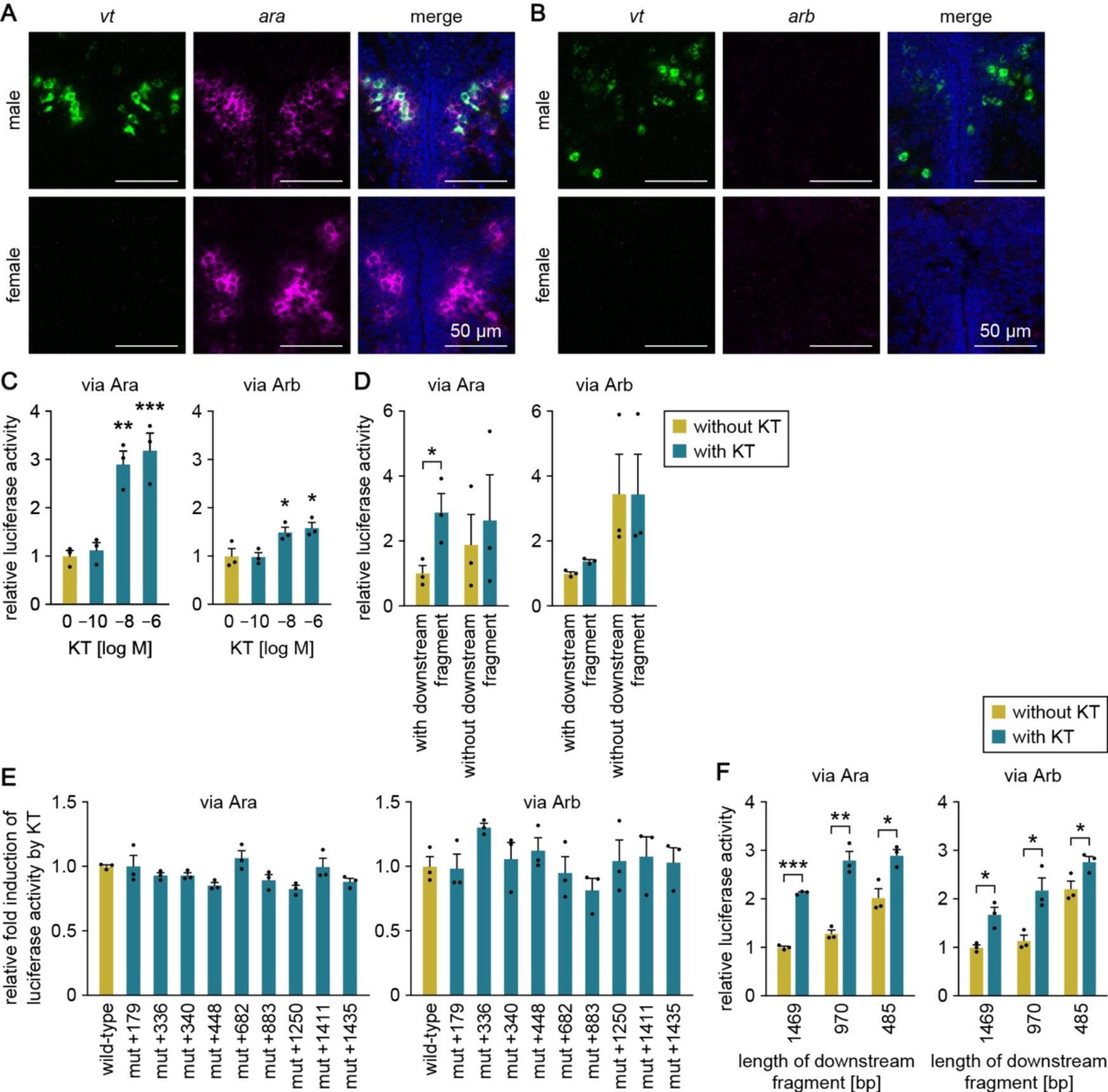
Androgens can directly activate the transcription of *vt* through Ara. (A, B) Representative images of the expression of *ara* (A) and *arb* (B) in the pNVT, where *vt* exhibits sexually dimorphic expression. In each row, left panels show *vt* expression (green), middle panels show *ara*/*arb* expression (magenta), and right panels show the merged images with DAPI staining (blue). Scale bars are all 50 μm. (C) Stimulation of *vt* transcriptional activity by KT. A luciferase-based reporter construct containing genomic fragments upstream of the first methionine codon and downstream of the stop codon of *vt* was transfected into COS-7 cells, in conjunction with an Ara or Arb expression construct. Cells were stimulated to varying concentrations of KT, and luciferase activity was measured. Fold induction was calculated relative to unstimulated cells. (D) Effects of removing the downstream fragment on KT-induced luciferase activity. A luciferase reporter construct with or without the downstream fragment was transfected into cells, along with the Ara or Arb expression construct. Cells were stimulated with or without KT (blue and yellow columns, respectively), and luciferase activity was determined. Fold induction was calculated relative to the construct with the downstream fragment without KT stimulation. (E) Effects of mutations in ARE-like sequences on KT-induced luciferase activity. A wild-type luciferase reporter construct (wild-type) or a construct carrying a mutation in the ARE-like sequence at position +179 (mut+179), +336 (mut+336), +340 (mut+340), +448 (mut+448), +682 (mut+682), +883 (mut+883), +1250 (mut+1250), +1411 (mut+1411), or +1435 (mut+1435) was transfected into cells, along with the Ara or Arb expression construct. Cells were stimulated with or without KT, and fold induction of luciferase activity was calculated relative to the wild-type construct. (F) Effects of 3′-truncation of the downstream fragment on KT-induced luciferase activity. Cells were transfected with a reporter construct containing 1469-bp, 970-bp, or 485-bp downstream fragment and an Ara or Arb expression construct. Cells were stimulated with or without KT (blue and yellow columns, respectively), and fold induction of luciferase activity was calculated relative to the construct containing the 1469 bp fragment without KT stimulation. Statistical differences were determined by Dunnett’s *post hoc* test (versus unstimulated control (C) or wild-type construct (E)) and unpaired t-test (D and F). \**p* < 0.05; \*\**p* < 0.01; \*\*\**p* < 0.001.

Next, we tested the ability of androgens to directly activate the transcription of *vt*. Our search for potential AREs in the *vt* locus of medaka identified six bipartite ARE-like sequences within the upstream flanking region (positions −2600, −2225, −2168, −1620, −817, and −242 relative to the first methionine codon) and nine within the downstream flanking region (positions +179, +336, +340, +448, +682, +883, +1250, +1411, and +1435 relative to the stop codon) (Supplementary Fig. 2). No ARE-like sequences were found in the gene body of *vt*. A transcriptional activity assay using a luciferase-based reporter construct containing the upstream and downstream flanking fragments of *vt* that carried these ARE-like sequences revealed that luciferase activity was dose-dependently induced by KT in the presence of either Ara (*p* = 0.0015 at 10^-8^ M and 0.0006 at 10^-6^ M) or Arb (*p* = 0.0437 at 10^-8^ M and 0.0201 at 10^-6^ M), with higher induction observed for Ara (Fig. 2C). An additional assay using only the upstream fragment (the downstream fragment was removed from the reporter construct) resulted in loss of KT-induced luciferase activity in the presence of Ara (*p* = 0.0379 and 0.6724 with and without the downstream fragment, respectively), suggesting that the *cis*-element responsible for KT induction is present in the downstream region (Fig. 2D). To further explore this finding, we introduced point mutations in each of the nine ARE-like sequences located in the downstream fragment in the reporter construct and studied the resulting changes in luciferase activity. However, none of the mutations had a significant impact on KT-induced luciferase activity in the presence of either Ara or Arb (Fig. 2E).

We therefore performed additional assays using reporter constructs containing serially 3′-truncated downstream fragments to examine the effects of simultaneous loss of multiple ARE-like sequences. However, even after truncating the downstream fragment from 1469 bp to 485 bp, KT-induced luciferase activity was not abolished in the presence of either Ara (*p* < 0.0001, = 0.0015, and 0.0178 for 1469, 970, and 485 bp fragments, respectively) or Arb (*p* = 0.0136, 0.0198, and 0.0487, respectively). These results suggest that the 485-bp region downstream of the stop codon of *vt* contains the *cis*-element responsible for KT induction. Considering that none of the mutations in the ARE-like sequences found in this region abolished KT induction, it may be that AR interacts with *cis*-elements other than the canonical ARE to activate the transcription of *vt*. Overall, these findings indicate that androgens can act on *vt*-expressing neurons in the pNVT to directly stimulate the transcription of *vt* via Ara, although the *cis*-element responsible for this process remains to be identified.

### Administration of exogenous Vt elicits aggression in females

If the high levels of aggression typically observed in male medaka result from male-specific production of Vt in the pNVT, then female medaka should also exhibit aggression when given exogenous Vt. We tested this idea by treating females with Vt and analyzing the resulting changes in intrasexual aggression. Aggression in medaka and many other teleosts includes five behavioral acts: chase, fin display, circle, strike, and bite (Oliveira *et al*., 2011; Kagawa, 2013). Vt treatment led to a significant increase in the number of chases (*p* = 0.0086), fin displays (*p* = 0.0104), and bites (*p* = 0.0011), but no strikes were induced (Fig. 3). Although not statistically significant, the treatment also evoked circles, which are not typically observed in females (Fig. 3). From these findings, it was evident that exogenous Vt elicits aggression in female medaka.

**Fig. 3.**
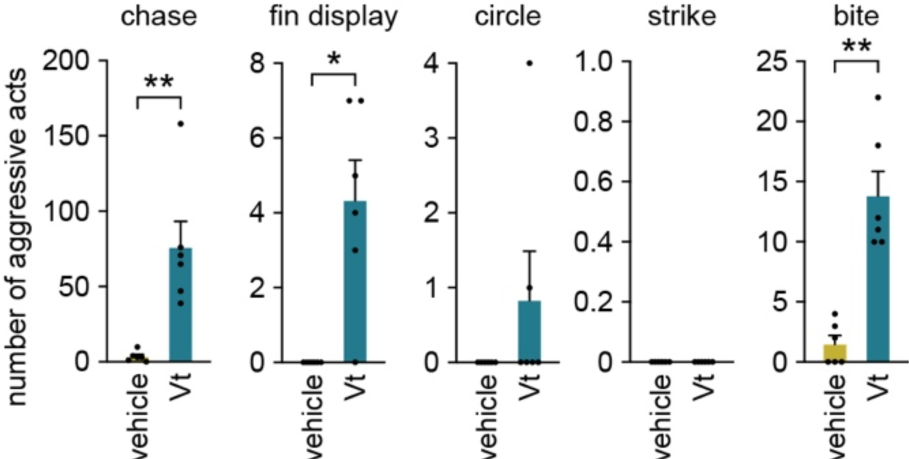
Administration of exogenous Vt elicits aggression in females. Shown is the sum of each aggressive act (chase, fin display, circle, strike, and bite) exhibited by females receiving vehicle only or Vt (n = 6 for each treatment). Statistical differences were determined by unpaired *t*-test with Welch’s correction. \**p* < 0.05; \*\**p* < 0.01.

### Altering the androgen milieu leads to correlated changes in the levels of aggression and *vt* expression in the tuberal hypothalamus

If androgen-dependent *vt* expression in the pNVT facilitates aggression, there is likely to be a correlation among the levels of *vt* expression in the pNVT, androgen action, and aggression. To explore this relationship, we administered KT to females with intact ovaries and evaluated changes in intrasexual aggression and *vt* expression in the pNVT. There was a significant increase in the number of chases after 4 days of KT treatment (*p* = 0.0421), and this increase persisted throughout the remaining treatment period (*p* = 0.0136 on day 5 and 0.0054 on day 7) (Fig. 4A). Although not statistically significant, the number of fin displays, circles, strikes, and bites also increased (Fig. 4A). *In situ* hybridization analysis revealed that KT treatment, even without ovariectomy, induced *vt* expression in the pNVT (*p* = 0.0043), along with increased aggression (Fig. 4B and C). In other brain nuclei, no significant changes in *vt* expression were noted with KT treatment (Fig. 4B).

**Fig. 4.**
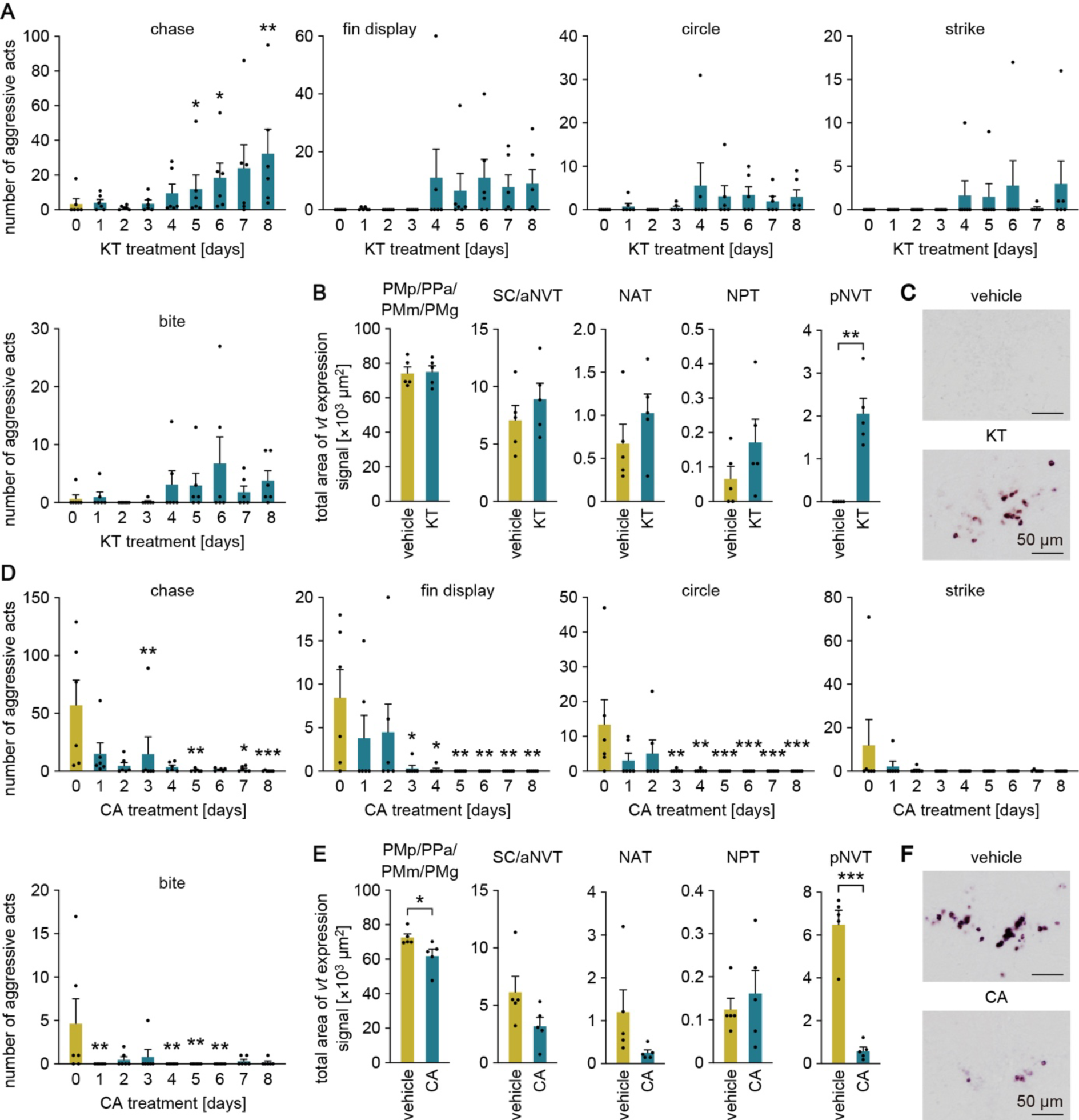
Altering the androgen milieu leads to correlated changes in the levels of aggression and *vt* expression in the tuberal hypothalamus. (A) Daily changes in each aggressive act (chase, fin display, circle, strike, and bite) exhibited by KT-treated females (n = 6). (B) Total area of *vt* expression signal in each brain nucleus of vehicle- or KT-treated females (n = 5 per treatment). (C) Representative images of *vt* expression in the pNVT of vehicle- or KT-treated females. Scale bars are 50 μm. (D) Daily changes in each aggressive act (chase, fin display, circle, strike, and bite) exhibited by males treated with the androgen receptor antagonist CA (n = 6). (E) Total area of *vt* expression signal in each brain nucleus of vehicle- or CA-treated males (n = 5 per treatment). (F) Representative images of *vt* expression in the pNVT of vehicle- or CA-treated males. Scale bars are 50 μm. Statistical differences were determined by Dunnett’s *post hoc* test (versus day 0) (A, D), except for the fin display data in panel D, which was determined by Dunn’s *post hoc* test, and unpaired *t*-test with or without Welch’s correction (B, E). \**p* < 0.05; \*\**p* < 0.01; \*\*\**p* < 0.001.

Next, we treated males with intact testes with the AR antagonist CA to inhibit androgen/AR signaling and conducted similar behavioral and expression analyses. The number of chases, fin displays, and circles was significantly reduced after 3 days of treatment (*p* = 0.0094, 0.0044, 0.0289, and 0.0007 for chases on days 3, 5, 7, and 8, respectively; *p* = 0.0323, 0.0275, and 0.0038 for fin displays on days 3, 4, and 5 onward, respectively; *p* = 0.0024, 0.0024, and 0.0003 for circles on days 3, 4, and 5 onward, respectively), and the number of bites was significantly reduced after 1 day of treatment (*p* = 0.0075 on days 1, 4, 5, and 6) (Fig. 4D). *In situ* hybridization analysis revealed that CA treatment substantially attenuated *vt* expression in the pNVT (*p* = 0.0005) (Fig. 4E and F). There were no notable changes in *vt* expression in other brain nuclei, except for a slight decrease in the PMp/PPa/PMm/PMg (*p* = 0.0402) (Fig. 4E).

Taken together, these results demonstrate a clear correlation among *vt* expression in the pNVT, androgen action, and aggression, and strengthen our hypothesis that the high levels of aggression typical of males are attributable to androgen-dependent *vt* expression in the pNVT.

### *vt* deficiency fails to prevent androgen-induced aggression in females

Lastly, we further tested our hypothesis by producing a *vt* knockout medaka line and comparing the aggression levels of females treated with KT between genotypes. We considered that, if our hypothesis is correct, KT treatment should elicit aggressive behavior in *vt*^+/+^ and *vt*^+/−^ females, but not in *vt*^−/−^ females. We employed TALEN-mediated genome editing to produce a medaka line with a deleterious frameshift mutation in *vt* (Supplementary Fig. 1A), resulting in an inability to produce the mature Vt peptide (Supplementary Fig. 1B). We then treated *vt*^+/+^, *vt*^+/−^, and *vt*^−/−^ females with KT and analyzed their aggression levels on days 0 (before treatment), 5, and 9. Contrary to our expectations, however, aggression was induced even in *vt*^−/−^ females, and no significant differences between genotypes were found for any aggressive act (Supplementary Fig. 1C). Therefore, this analysis did not support our hypothesis.

## Discussion

We previously identified male-specific populations of VT-expressing neurons in the tuberal hypothalamus of medaka (Kawabata *et al*., 2012). Here we proposed and tested the hypothesis that VT expression in these neuronal populations is induced exclusively in males in an androgen-dependent manner and contributes to high, male-typical levels of aggression. Teleost fish, like terrestrial vertebrates, have a major population of VT-expressing neurons in the brain nucleus homologous to the PVN and its surrounding areas (PMp/PPa/PMm/PMg) (Godwin and Thompson, 2012). It has been reported in several teleost species, including medaka, that the number of one or more subpopulations of these VT-expressing neurons correlates with the level of aggression, and for this reason, male-typical aggression in teleosts has been attributed to this neuronal population (Godwin and Thompson, 2012; Kagawa, 2013; Silva and Pandolfi, 2019). On the other hand, no attempt has been made to assess the contribution of the tuberal hypothalamic populations to aggression.

Here we first demonstrated that male-specific VT expression in the tuberal hypothalamus is indeed dependent on androgens. More specifically, VT expression in the pNVT of the tuberal hypothalamus is induced exclusively in post-pubertal males by large amounts of androgens secreted by the testes. To our knowledge, this is the first report showing that teleosts, although lacking the VT-expressing neurons in the extended amygdala that are androgen-dependent and thus male-biased, have VT-expressing neurons with comparable properties in a distinct brain nucleus. We observed no sex steroid dependence of VT expression in other brain nuclei including the PMp/PPa/PMm/PMg, as reported in other teleost species such as bluehead wrasses (*Thalassoma bifasciatum*) (Semsar and Godwin, 2003). In teleosts, therefore, the neuronal population in the pNVT is presumably responsible for reproductive cycle-dependent VT functions, including increased aggression during the mating period.

Despite the similarities, there are important differences in androgen-dependent VT expression in the extended amygdala of terrestrial vertebrates and the tuberal hypothalamus of teleosts. First, in the rodent extended amygdala, a large fraction of testosterone is converted locally to E2 and then acts through the ESR (Aspesi and Choleris, 2022), whereas in the medaka tuberal hypothalamus, KT, an androgen that cannot be converted to E2, acts directly through the AR. Because the effects of androgens on male behavior rely primarily on an AR-mediated pathway in many vertebrate species (Okubo *et al*., 2022), our findings in medaka may be broadly applicable to other species. The second difference concerns whether treating adult females with sex steroids induces VT expression above basal levels. In adult female rats, VT expression in the BNST is elevated by estrogen treatment but only to levels observed prior to ovariectomy (De Vries *et al*., 1994; Turano *et al*., 2019). This is because, in rodents, and probably in other terrestrial vertebrates, sex steroids produced in the fetal gonads in a sex-specific manner act on the developing BNST to shape irreversible and enduring sex differences in VT expression (Rigney *et al*., 2022; 2023). Furthermore, in rodents, this sex difference is due in part to the direct neuronal actions of sex chromosome-linked genes, which are independent of sex steroid effects (De Vries *et al*., 2002). In medaka, by contrast, treating females with androgens markedly induced VT expression in the pNVT, which was virtually absent in ovary-intact and ovariectomized conditions. Taken together with the observation that castration in males severely reduced VT expression in the pNVT, it seems that the sexually dimorphic pattern of VT expression in medaka can be reversed between the sexes in response to changes in the adult androgen milieu. It is known that the sexual phenotypes of teleosts (behavioral or otherwise), unlike those of terrestrial vertebrates, are highly labile across the lifespan and can be reversed between the sexes, even as adults (Okubo *et al*., 2019; 2022); the present study now shows that manipulation of the adult androgen milieu effectively reverses sex-typical aggressive behavior. The reversibility of sexually dimorphic VT expression may reflect this fact.

It may be relevant to note here that KT-induced *vt* expression in the pNVT of females was apparent but at a relatively low level compared to that of castrated males. A possible explanation for this observation is that full induction of *vt* expression by KT in females requires a relatively long period of time because it involves the generation of new neurons or the activation of the counterparts of male *vt*-expressing neurons that are in a quiescent, non-activated state. It would be worthwhile to determine if treating females with KT for a longer period of time results in comparable levels of *vt* expression as males.

The question then arises whether androgen/AR signaling induces VT expression in pNVT neurons directly or indirectly through other target genes or cells. We found that *ara*, one of the two AR genes in teleosts, is expressed in almost all VT-expressing neurons in the pNVT. We further showed by transcriptional activity assays that androgens can directly stimulate the transcription of *vt* and that the magnitude of this stimulation is greater through Ara than through Arb. These results suggest that androgens directly activate VT expression in the pNVT through binding to Ara. This finding is consistent with recent work in cichlids (*Astatotilapia burtoni*) showing that *ara* and *arb* are functionally differentiated, with *ara* responsible for promoting aggression (Alward *et al*., 2020). Although it is not known whether VT is a direct transcriptional target of AR signaling (or ESR signaling) in terrestrial vertebrates, as it is in medaka, androgen-induced *Vt* expression in the BNST of adult rats has been reported to involve changes in the DNA methylation pattern of the *Vt* promoter (Auger *et al*., 2011). Future work will be needed to clarify whether a similar mechanism exists in the medaka pNVT and to identify functional AREs in the medaka *vt* locus (which eluded detection in the present survey) in order to clarify the evolutionary conservation and divergence of regulatory mechanisms for sexually dimorphic VT expression.

We also tested the hypothesized role of male-specific VT expression on aggression. Although VT has been implicated in aggressive behavior in many teleost species (Godwin and Thompson, 2012; Silva and Pandolfi, 2019), Yokoi *et al*. (2015) observed no anomalies in intrasexual aggression of male medaka carrying a missense mutation in *vt* (leading to replacement of the first methionine with arginine and loss of the ability to produce the mature Vt peptide). While this observation may suggest that VT is not involved in aggression in medaka, we showed here that administration of exogenous Vt peptide elicited aggressive behavior in female medaka, which typically exhibit little or no aggression. This result indicates that VT certainly serves to facilitate aggression in medaka as well as in other teleosts (although it is possible that the induced aggression may represent a pharmacological effect rather than a physiological role of VT). Furthermore, manipulation of the androgen milieu of adult male and female medaka to inhibit or facilitate aggression revealed a clear correlation among *vt* expression in the pNVT, androgen action, and aggression. More specifically, treatment of male medaka with an AR antagonist resulted in a marked reduction in both aggression and *vt* expression in the pNVT, whereas androgen treatment of females elicited both of them. All of these findings support our hypothesis that male-specific VT expression contributes to male-typical high levels of aggression, thereby strengthening its validity.

Finally, we further tested this hypothesis using *vt*-deficient female medaka but, contrary to our expectations, androgen treatment induced comparable levels of aggression in *vt*-deficient females as in wild-type females. A straightforward interpretation of this result would suggest that our hypothesis is incorrect and androgen-induced aggression is not mediated by VT; however, the lack of detectable effects of *vt* deficiency might also be due to functional compensation by other genes or pathways. A good candidate for this functional compensation is OT, a nonapeptide closely related to VT that can partially activate VT receptors (Mennigen *et al*., 2022; Rae *et al*., 2022). In medaka, *ot*-expressing neurons are not observed in the pNVT, where male-specific *vt*-expressing neurons reside (Kawabata *et al*., 2012), but it is possible that *ot*-expressing neurons in another brain nucleus project to the same brain region as male-specific *vt*-expressing neurons and compensate for *vt* function. Contrary to male medaka carrying a missense mutation in *vt*, which showed comparable levels of aggression to wild-type males, as described above (Yokoi *et al*., 2015), male medaka carrying a missense mutation in the VT receptor subtype *v1a2* (leading to replacement of asparagine at position 68 with isoleucine) were found to be less aggressive (Yokoi *et al*., 2015). This discrepancy can also be explained if OT compensates for the loss of VT function via V1A2. It would be interesting to test this idea in future studies by producing double-knockout medaka for *vt* and *ot* and analyzing their behavioral phenotypes.

In summary, our results have demonstrated that VT expression in the tuberal hypothalamic nucleus is induced exclusively in males, most likely as a result of direct transcriptional activation of VT by androgen/Ara signaling, although it remains to be determined whether this VT expression is relevant to the high levels of aggression typical of males. The anterior hypothalamus of rodents, which is homologous to the teleost pNVT, contains no VT-expressing neurons; however, it receives heavy projections from male-biased VT neurons in the BNST and abundantly expresses the behaviorally-relevant V1A receptor, thus representing a primary site of action of VT for aggression (Rigney *et al*., 2022; 2023). If the V1A receptor homolog is also expressed in the teleost pNVT, VT produced in this brain nucleus may contribute to male-typical aggression by acting in an autocrine/paracrine manner. Further studies, including analyses of receptor expression and axonal projection patterns, will be needed to test this idea. It would also be worthwhile investigating whether the current findings in medaka apply to other teleosts. The resulting information on species variation in the spatiotemporal expression patterns and regulatory mechanisms of VT may provide some insight into the diversity of social structures and behavioral patterns in teleosts.

## Acknowledgments

The authors would like to thank the National BioResource Project (NBRP) Medaka for providing the BAC clone, Dr. Junpei Yamashita and Takayasu Tsumaki for technical assistance, and Dr. Towako Hiraki-Kajiyama and Akio Takeuchi for experimental support. This work was supported by the Ministry of Education, Culture, Sports, Science, and Technology (MEXT), Japan, and the Japan Society for the Promotion of Science (JSPS) (MEXT/JSPS grant numbers 13J04816 (to YKS), 17H06429, and 23H02305 (to KO)).

## Declarations of interest

The authors declare no competing interests.

## Notes

### Competing Interest Statement

The authors have declared no competing interest.

