## Supplementary Materials for "Male-specific vasotocin expression in the medaka tuberal hypothalamus: androgen dependence and probable role in aggression"

**Supplementary Table 1. Abbreviations of medaka brain regions and nuclei.**

| abbreviation | full name | location |
|---|---|---|
| brain region | | |
| Cb | cerebellum | |
| Hyp | hypothalamus | |
| MO | medulla oblongata | |
| OB | olfactory bulb | |
| OpN | optic nerve | |
| OT | optic tectum | |
| Tel | telencephalon | |
| brain nucleus | | |
| aNVT | anterior part of the ventral tuberal nucleus | hypothalamus |
| NAT | anterior tuberal nucleus | hypothalamus |
| NPT | posterior tuberal nucleus | hypothalamus |
| PMg | gigantocellular portion of the magnocellular preoptic nucleus | preoptic area |
| PMm | magnocellular portion of the magnocellular preoptic nucleus | preoptic area |
| PMp | parvocellular portion of the magnocellular preoptic nucleus | preoptic area |
| pNVT | posterior part of the ventral tuberal nucleus | hypothalamus |
| PPa | anterior parvocellular preoptic nucleus | preoptic area |
| SC | suprachiasmatic nucleus | hypothalamus |

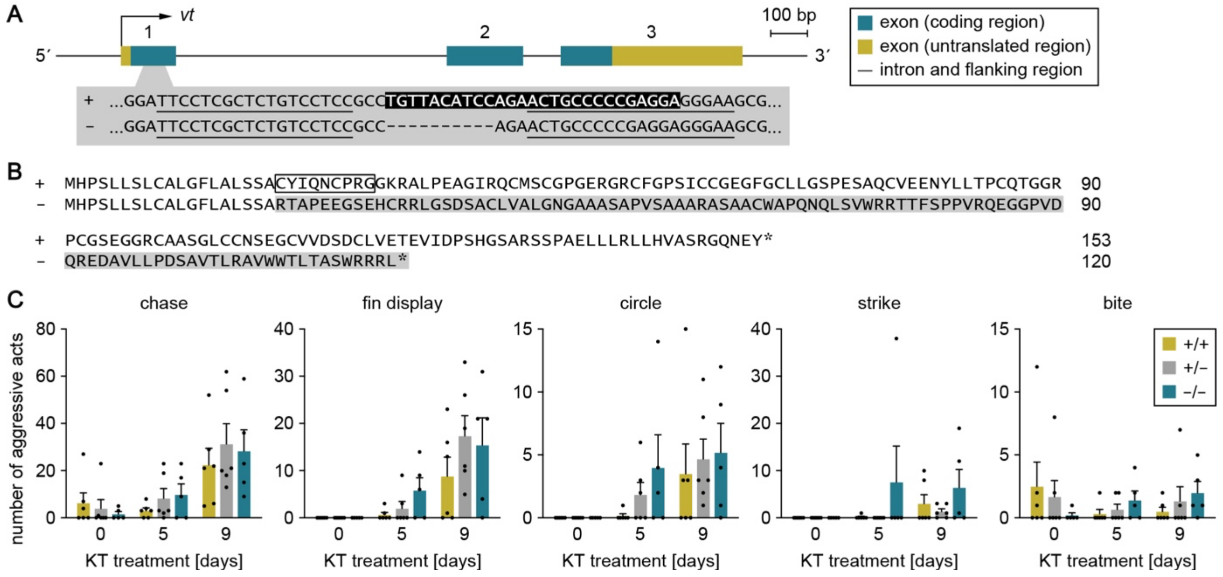


**Supplementary Fig. 1. *vt* deficiency fails to prevent androgen-induced aggression in females.** The *vt* knockout medaka line was produced using TALEN technology. (A) Schematic of the gene structure of *vt* and the relative position of the target site. Each exon is numbered above. Nucleotide sequences of the wild-type (+) and mutant (−) alleles at the target site are shown. The mature Vt peptide-coding sequence is highlighted by white letters on a black background. Deleted nucleotides are marked with dashes, and TALEN-binding sites are underlined. (B) Comparison of the amino acid sequences deduced from the + and − alleles. The sequence altered by the frameshift is shaded in gray. The mature Vt peptide region is represented by a box. Asterisks denote stop codons. (C) Changes in each aggressive act (chase, fin display, circle, strike, and bite) exhibited by $vt^{+/+}$ (n = 6), $vt^{+/-}$ (n = 6), and $vt^{-/-}$ (n = 5) females treated with KT. Statistical differences were determined by Bonferroni's *post hoc* test (C).

GGAGGTTATAGACCCATCCCATGGCTCTGCCAGAAGCTCGCCTGCAGAGCTGCTGCTGCGTCTCCTACACGTCGCCAGCAGAGGACAGAACGAATACTGA −1

AACCTGGATCTGCAGCAGACACGACGCCTCAACTTCCTCTCAGCCAAATGAGACATTGCTCTTTGCTCTCTGCTTGAACTCTCCCCTTTTAGGATACTGG 100

CATACATATCCAGGGGAAAAAATACTCGTAGATTTCATCTTTGTTCCCTCTTAAAGAAATGTACATATGACTGCACATATTGTAAACAATTCAGAGCCTC 200

TGTTCAAATAGATAAGAGCTAACAGCAAACTGTAAATAAGTCACAGATTGAAATTTAAAAATAGATGCTCATAACAATTACGAAATGGCTCTCATGTAGT 300

AGAACTATGCCTGTAGATTGAAAACAAGAGAAAATACAGAATGTCTCATCACAGCCAAAACGCCTTCTGCTGTTTTTCTTTATCTGTAGCCCCAAACTAA 400

GAAGACTTATTTTACAGGATGGTGAGAATTTGTGTGAAAAGGAACCAAATGTAATTTGAGCGATCTCAGAGATGAGTAATCATCCACAACAATTGCAAAT 500

GTCCCTCTTTAGGTGGTCTTTCTTTCCTTCCAATATTTTGACATGTCTTGCTCCTGATGCACCTCTCTACCAGCTGTGTATCAACATGTCACGGTCTGTT 600

AATCAAAGGCAATGGCATTTCATTTGAGGGATTGGGCTTTGTGTGCGTGCGATGACTGAAGGCTGGGTTACGTGCCAACAATGAGAGGAAAGACGGTTTG 700

TCTTTTAAAGTCTGTTTCAACCTGGAGGTTGACTGCAAGGCACCGCCTTGTACTTGACAGTTACATGTTTTGACTGATAGAGCTTGGATCTTTTGCAAAG 800

GCTGAACCTTAAATCCTGTTGATATGAGTGATAAATCACTGATAATCCCCAACTTGGAGCTTCTTCGGCCCCCTGACAACAACAAGAGCGGGATGTCCAC 900

ACACATTCATCTAAGGAATTTGATTGTTTTGTAATGTATAATCACTACAACGATTCTGGTAATCAAGTTCTGCTCAATGTTAACCATGTAAATAAAATTA 1000

CCTTAATAACCTTACTTCCTTTCAATGGACATATAATTTGGTGCAAAAAAAACTCTTTAAATTTGGAGATTCTACCTTTTTGCTATCAATGCGTTCAGAC 1100

TGAATATTTTACGTACTCTCAGAATACTTCTAAAGTCAAAGAAATACACAATATCACCAAAAGAATTCGCTCACCCATGATCAGAATCAGGTGTCCTGAT 1200

CCCTTGGTCTGGTCACAGGTGTATAAATCAAGCACTCAGGCATGCAGACTGTTTCTTACAACTATTTGTGAAAGAATGGGCCGCTCTCAGGGGTTCAGTG 1300

ATTTCTAGTGTGGAAACTGTCCCAGGATAGCACCTGTGCAACAAATCCAGTCATGAAATGTCCTCACCCCTTAAATATTCCGCAGTCAATTGTCAGCTCC 1400

ACAAGAACAAAATGGAAGAGTTTGGGAACAACAGCAACTCAGCCATAAAATTGAAGGCCTTGTAAACTTTCGGAGAGGGCTCCGTGGATGAGAAGCGCAT 1500

**Supplementary Fig. 2. Canonical bipartite ARE-like sequences in the downstream flanking region of *vt*.** Nucleotide sequences of the 3′-end of *vt* and its downstream flanking region are shown. The coding region is shaded in gray and the 3′-untranslated region is underlined. ARE-like sequences are indicated in yellow with the ARE half-sites boxed. Nucleotide numbers relative to the stop codon are shown at the right of each line.
